# Longevity-associated SMAD3 non-coding centenarian variant impairs a cell-type specific enhancer to reduce inflammation

**DOI:** 10.1101/2023.05.17.540984

**Authors:** Jiping Yang, Archana Tare, Lei Zhang, Seungsoo Kim, Seungjin Ryu, Qinghua Guo, Yizhou Zhu, Xizhe Wang, Xifan Wang, Adam Hudgins, Di Guan, Chen Jin, Hyun-Kyung Chang, Gil Atzmon, Sofiya Milman, Nir Barzilai, Jan Vijg, Laura Niedernhofer, Paul D. Robbins, Yousin Suh

## Abstract

Given the pro and anti-geronic roles of the TGF-β superfamily in aging, we hypothesized that human longevity involves genetic variation in TGF-β signaling genes. Here we utilized a candidate functional genomic approach to identify and characterize functional variants in TGF- β signaling associated with human longevity. Targeted sequencing of 113 genes involved in aging-associated TGF- β signaling in an Ashkenazi Jewish centenarian cohort identified genetic variants robustly associated with human longevity. In particular, a centenarian-enriched intronic variant residing in a cell-type specific enhancer in SMAD3, a critical receptor-regulated TGF- β signal transducer, was identified. This non-coding SMAD3 variant (rs8040709) altered binding of ELK1, a member of the ETS family of transcription factor important for enhancer activity in certain cell types, resulting in reduced SMAD3 expression. Analysis of the variant in cell types derived from gene edited iPSCs demonstrated the variant reduced SMAD3 expression, senescence and inflammation in endothelial cells. In addition, heterozygosity in SMAD3 improved healthspan and reduced senescence in the *Ercc1*^-/Δ^ progeroid mouse model of accelerated aging. Taken together, these experiments demonstrate that variants in a cell type specific enhancer of SMAD3 resulted in reduced expression, senescence and inflammation and contributes to human longevity. Thus, SMAD3 represents a validated targeted for drug development for extending human healthspan.

## Introduction

By 2050, the number of people older than 80 years is projected to triple globally, which would lead to an unprecedented increase in the burden of age-related diseases and healthcare costs. Aging is the major risk factor for many chronic diseases. Geroscience research hypothesizes that targeting the underlying mechanisms of aging, rather than its consequences, holds the potential to simultaneously delay the onset and progression of multiple age-related pathologies^1^. Results from heterochronic parabiosis experiments indicate that young circulation system contains factors responsible for maintenance of youthful characteristics and reversal age-related dysfunction in multiple organs throughout the body in old animals. So far, multiple geronic factors such as GDF11^2^, TIMP2^3^, B2M^4^, CCL11^4^ and osteocalcin^5^ have been identified as capable of recapitulating some of the effects of heterochronic parabiosis. Interestingly, most of the identified systemic geronic factors are either members of transforming growth factor beta (TGFβ) superfamily or are directly modulated by TGF-β signaling, which is one of the major pathways regulating secretory factors.

Centenarians represent the ideal human model to study longevity since they generally age slower and healthier due to a strong genetic component. Given the strong association of TGF- β signaling with both geronic factors and aging, we hypothesize that human longevity involves genetic variation in TGF- β signaling genes. Here, we apply a candidate functional genomic approach to identify and characterize functional variants associated with human longevity. Using targeted sequencing of 113 genes involved in aging-associated TGF- β signaling in an Ashkenazi Jewish centenarian cohort, we identified genetic variants robustly associated with human longevity. Function annotation was performed within a holistic framework incorporating genomic and epigenomic information to prioritize nominally significant variants. This approach identified a centenarian-enriched intronic variant residing in a cell-type specific novel enhancer in SMAD3, a critical receptor-regulated TGF- β signal transducer. Next, we comprehensively characterize the function of the non-coding SMAD3 variant (rs8040709) on SMAD3 gene expression and elucidate the underlying molecular mechanisms, including altered transcription factor binding that mediate the enhancer activity and cellular consequences of SMAD3 down- regulation on protection against senescence and inflammation. Moreover, we demonstrate that mimicking the impact of longevity-associated regulatory SMAD3 variant via heterozygous knock-out of Smad3 decreased senescence-associated secretory phenotypes (SASP) and extended healthspan in the Ercc1^-/Δ^ progeroid mouse model of accelerated aging.

## Results

### Sequencing of TGF-β signaling-related genes in an Ashkenazi Jewish centenarian cohort

We selected 113 candidate genes implicated in TGFβ signaling including (1) TGF-β superfamily members such as TGFBs, Growth and Differentiation Factors (GDFs), Bone Morphogenetic Proteins (BMPs), Myostatin, Activins/Inhibins and Nodals; (2) TGF-β regulators such as ligands, antagonists and inhibitors; (3) signal receptors and co-receptors; (4) signal transducer SMADs, SMAD regulators, transcriptional factors and transcriptional targets (Table S1). For 23 of the selected genes, we sequenced the whole gene body and 2 kb upstream from the transcription start site (TSS), except for GDF11, where 10 kb upstream was included. For the rest genes (namely exon covered), we included 2 kb upstream, all exons, and 20 bp of each exon-intron junction. The targeted regions totaled 2,199,107 bp with 95.3% base coverage.

From the sequencing of the targeted genomic regions in our Ashkenazi Jewish (AJ) centenarian cohort^6^, 37,616 variants were discovered and 93.7% of them were non-coding variants (Table S2, Fig. 1A and Fig. S1A). In order to identify longevity-associated variants, we performed variant based analysis using Fisher’s exact test. In general, rare variants tend to show higher significance compared to common variants (Fig. S1B). Also, vast majority of the most significant variants are rare or low-frequent variants. We further prioritized 95 significant variants with nominal p value less than 0.01 (Fig.1B).

**Figure 1.**
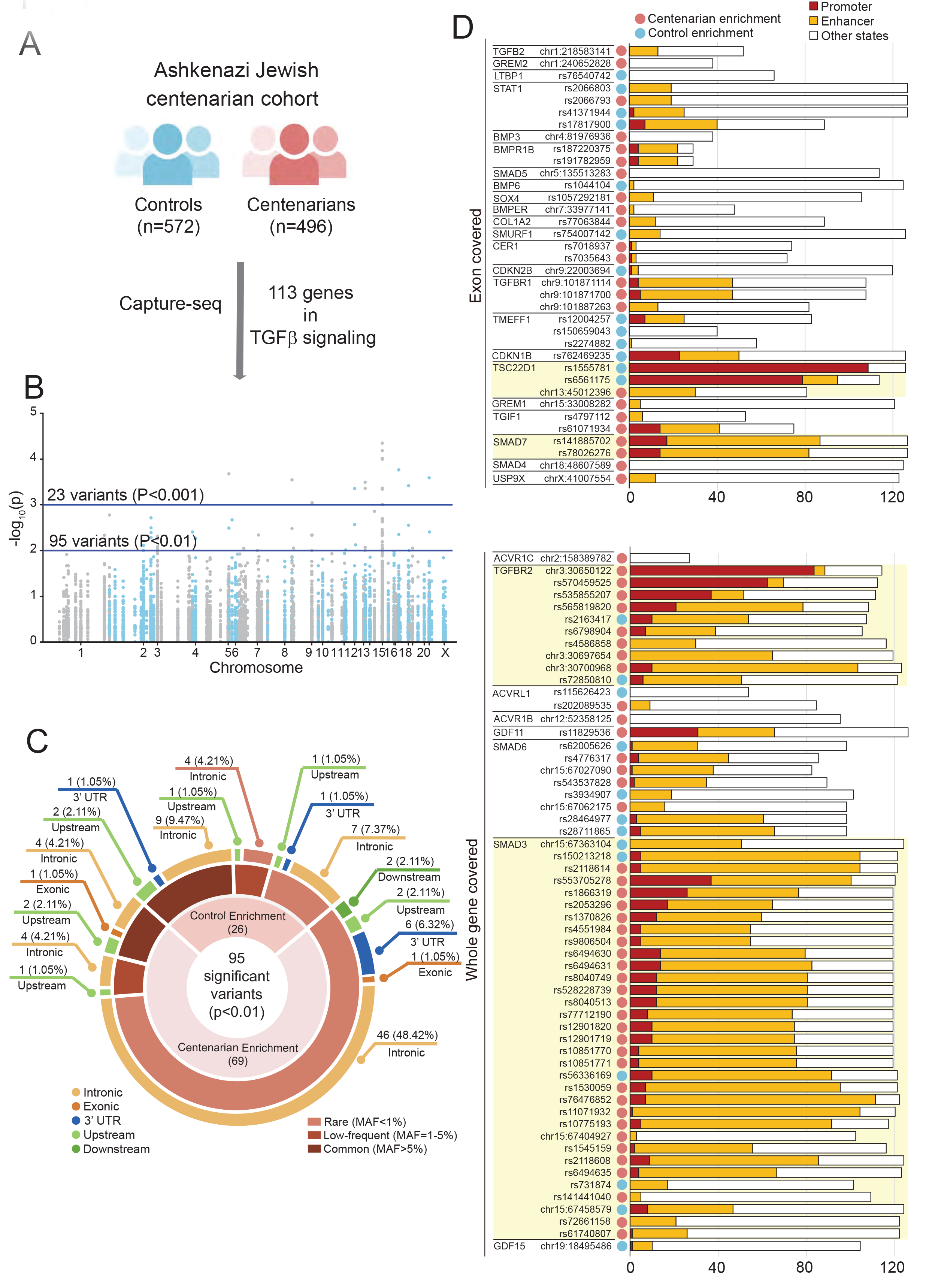
Capture-seq in an Ashkenazi Jewish centenarian cohort A. 113 TGF-β signaling-related genes were sequenced in an Ashkenazi Jewish centenarian cohort included 572 controls and 496 centenarian individuals. B. Manhattan plot showing all variants discovered in this longevity-genetic association study. C. 95 significant variants (P<0.01) categorized by allele frequency and region. D. Functional annotation of 93 non-coding variants by Roadmap Epigenomics. X- axis indicated the number of cell/tissue types showing epigenetic signatures (promoter in red, enhancer in yellow).

93 (97.89%) of them resided in non-coding regions (Fig.1C). Significant variants were enriched in centenarians, indicating that gain of protective variants instead of loss of deleterious variants are likely to contribute more to human longevity. Meanwhile, over 80% centenarian-enriched variants were rare variants, suggesting that rare variants have a stronger impact on human longevity. Meanwhile, 23 variants with nominal p value less than 0.001 exhibited similar features including 1) high proportion of non-coding variants; 2) more centenarian-enriched variants; and 3) all rare variants showed centenarian enrichment with a higher minor allele frequency in centenarian group (Fig. S1C).

Next, we focused on the 93 non-coding variants with nominal p value less than 0.01. To identify potential functional regulatory variants, we integrated Roadmap Epigenomics^7^ data including over 120 reference epigenomes from diverse cell types or tissues. 85 of the longevity-associated non-coding variants overlapped with cis-regulatory elements in at least one reference epigenome (Fig. 1D). At the gene level, *TSC22D1*, *SMAD7*, *TGFBR2* and *SMAD3* showed the enrichment of putative regulatory variants in over 50 tissues and cell types. Strikingly, almost one third of the regulatory variants were in the *SMAD3* gene (Fig. 1D). Although the sequencing coverage of SMAD3 was the longest in our capture-seq, the variant discovery rate (significant variant number/sequenced kb length) of SMAD3 was the highest among all gene body-covered genes and was still top when compared with exon-covered genes (Fig. S1D). This result strongly suggests that the *SMAD3* gene loci is a hotspot for rare variants associated with human longevity.

### Identification of a rare longevity-associated SMAD3 haplotype that lied in a cell-type specific enhancer in intron 1 of SMAD3

Since a group of SMAD3 rare variants in intron 1 showed centenarian enrichment, similar minor allele counts and location proximity, we checked the linkage disequilibrium (LD) of the variant set and identified a haplotype block containing 23 perfectly linked (R^2^=1, D’=1) variants (Table S3, Fig. 2A and 2B). None of these variants were in LD with exonic or splice variants. Due to the limitation of sample size, the integrated allele frequency information from AJ population (n=1,736) in gnomAD v3^8^ was examined. Minor allele frequency of the variants in this haplotype block in AJ centenarian was around 7-fold when compared with AJ population in gnomAD (p = 0.0009) (Table S3), confirming longevity-association of the rare haplotype.

**Figure 2.**
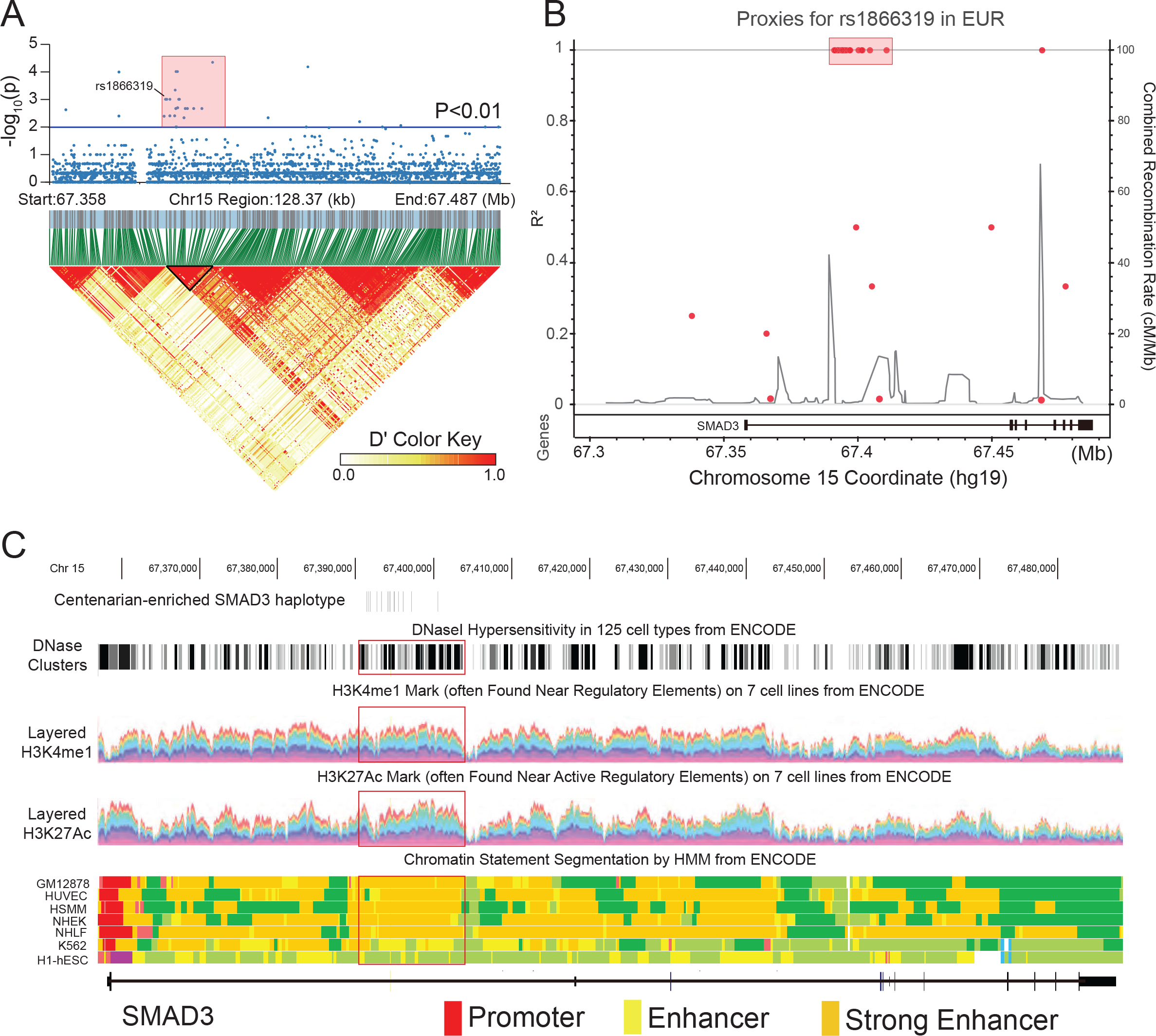
Identification of a longevity-associated haplotype in SMAD3 intron 1. A. Haplotype block in SMAD3 genes generated by variant data from 1000 Genomes Project. Most of the longevity-associated SMAD3 variants lie in a LD block (black rectangle and red highlighted). The first variant rs1866319 within this LD block was indicated. B. The linked variants of the lead variant rs1866319 was shown by LDproxy Tool using data of European population. C. Genome browser showing the DNase hypersensitivity, H3K4me1, H3K27Ac and ChromHMM information in *SMAD3* gene locus. Red square indicated the longevity-associated SMAD3 haplotype.

To predict the function of this longevity-associated SMAD3 haplotype, we scanned publicly available databases to determine if this haplotype was within potentially functional gene regulatory elements. The longevity-associated haplotype overlapped with an enhancer region with robust H3K4me1 and H3K27Ac marks according to ChromHMM^9^, a simple database characterizing the chromatin state in different cell types (Fig. 2C). This region showed strong enhancer signals in 5 cell types including GM12878, HUVEC, HSMM, NHEK and NHLF with weak or no enhancer signals in K562 and H1- embryonic stem cells (ESC). Next, we screened H3K27Ac ChIP-seq data from Encyclopedia of DNA Elements (ENCODE)^10^ and found this enhancer showed a cell-type/lineage-specific activity. In consistent with the findings in ChromHMM, this enhancer was inactivated in pluripotent stem cells including ESCs and induced pluripotent stem cells (iPSCs) (Fig. S2A). However, this enhancer was activated when ESCs were differentiated into the mesoderm lineage, but not into ectoderm and endoderm lineages *in vitro*. Similar to PSCs, this enhancer was silenced in brain tissues, brain-isolated cell derivates and hESC-differentiated neural progenitor cells and neurons (Fig. S2A). However, this enhancer exhibited strong activities in cells involved in hematopoiesis lineage including hematopoietic stem cells (HSC), B cells, T cells, monocytes, natural killer cells and neutrophils (Fig. S2A). In addition, active enhancers were observed in multiple mesoderm-lineage cells such as endothelial cells, mesenchymal stem cells and muscle cells as well as some fibroblast lines (Fig. S2A). Except for the enhancer marks, we also confirmed the chromatin interaction between the longevity-associated SMAD3 haplotype region and SMAD3 promoter in a HiC analysis of human blood T cells^11^ (Fig. S2B). Taken together, we identified a rare longevity-associated SMAD3 haplotype that lies in a cell-type specific enhancer in intron 1 of SMAD3.

### rs8040749 variant within the longevity-associated SMAD3 haplotype was the potential causal variant

Due to the linkage disequilibrium (LD) effect in the genome, identification of truly causal variant within a haplotype is challenging especially for non-coding variants in enhancer region. We looked up these variants on the RegulomeDB^12^ database, which scores variants based on experimental data from Encyclopedia of DNA Elements (ENCODE), Gene Expression Omnibus (GEO), and published literature. The SMAD3 rare haplotype contains 3 variants (rs8040749, rs10851770 and rs1530059) with a score of 2b which means likely to be functional. We hypothesized that the alteration of enhancer-promoter contacts requires the involvement of chromatin binding protein such as transcription factors. Therefore, whether the 3 potential functional regulatory variants can alter protein binding profiles was investigated by searching JASPAR^13^, a database of transcription factor binding profiles. Two variants (rs8040749 and rs108517708) were found to be located in conserved protein binding motifs (Fig. S3A). In particular, rs8040749 was located at a perfectly conserved site of canonical binding motif of ETS (E-Twenty-Six) family genes and centenarian-enriched G-allele created the binding site. In contrast, it was not crucial for NR6A1 binding since its position in binding motif was poorly conserved (Fig. 3A and S3B). The rs10851770 variant within ETS family genes binding motif may not to be critical to protein binding since both C and T allele exist naturally (Fig. 3A). Consistent with our prediction, we found ChIP-seq peaks of two ETS family genes ELK1 and ELF1 at rs8040749 locus, while no ChIP-seq peak was observed at rs10851770 locus from the ENCODE project (Fig. 3B). Taken together, we prioritized the SMAD3 variant rs8040749 as the potential causal variant.

**Figure 3.**
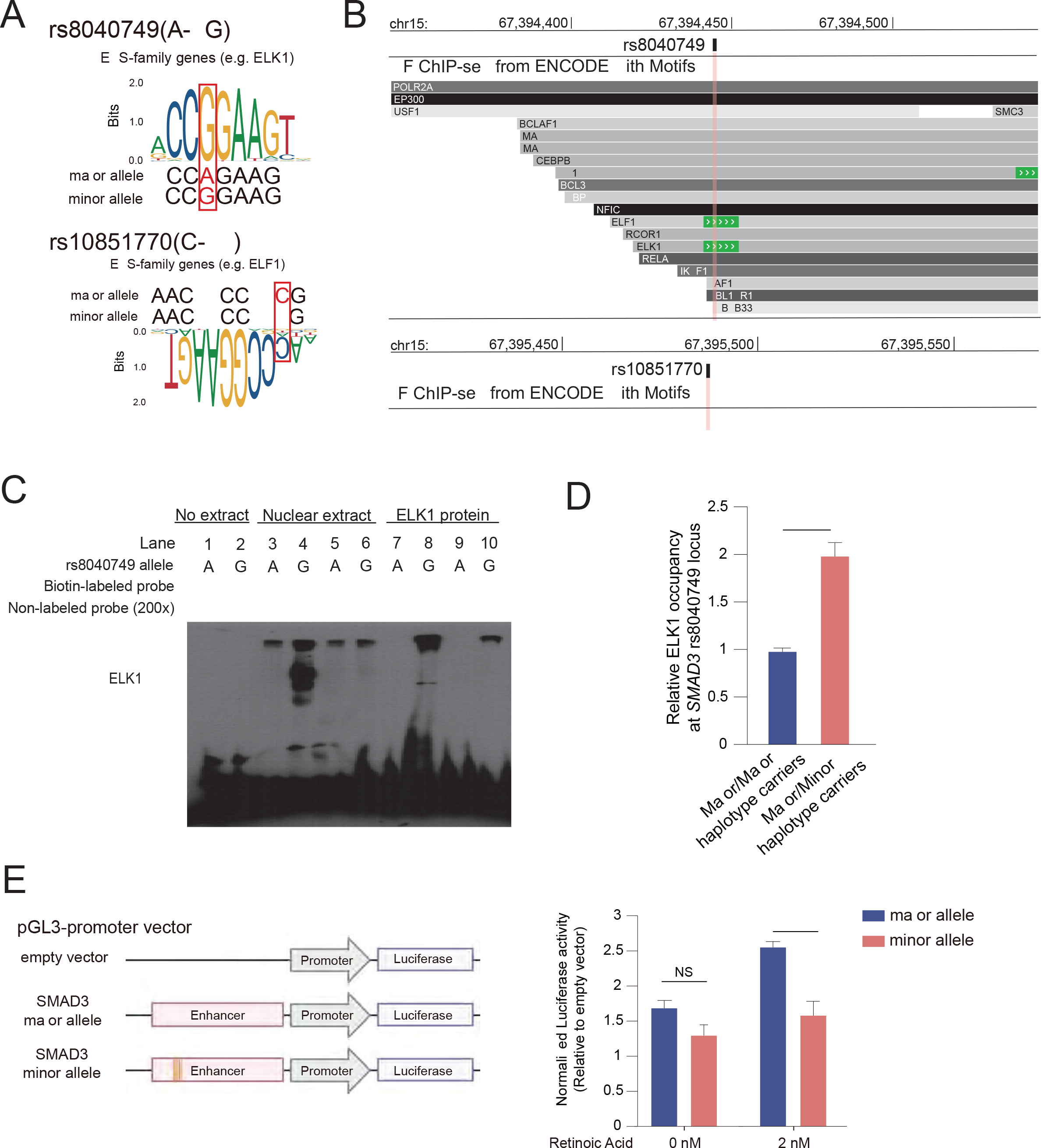
**Identification of rs8040749 as the causal variant** A. rs8040749 and rs10851770 overlapped with the binding site of ETS family genes, and the centenarian rs8040749-G allele was predicted to create a binding site. B. Genome browser showing the TF ChIP-seq from ENCODE with motifs at rs8040749 and rs10851770 loci. C. EMSA showing a stronger binding of ELK1 protein to rs8040749-G allele compared with rs8040749-A allele. D. ELK1 ChIP-qPCR in lymphoblastoid cell lines derived from major/major haplotype carriers (n = 3) and major/minor haplotype carriers (n = 3). **, p < 0.01. E. Dual luciferase reporter assays comparing enhancer activities of control and centenarian haplotype with or without treatment of Retinoic Acid. NS, not significant; **, p < 0.01.

To determine the protein binding specificity of each allele of rs8040749, we performed an Electrophoretic Mobility Shift Assay (EMSA) using allele-specific 20bp biotinylated double stranded oligos, nuclear extracts from LCLs, and recombinant ELK1 protein, and found that only the longevity-associated G-allele of rs8040749, not the reference A allele, was bound strongly by ELK1. A 200-fold molar excess of unlabeled probes outcompeted ELK1 binding at the G allele, but had no effect on the nonspecific signal at the A allele (Fig. 3C). To confirm the allele-specific binding of ELK1 in LCLs, we performed allele-specific chromatin immunoprecipitation (ChIP) TaqMan assay in longevity-associated haplotype heterozygous carrier LCLs. We detected a 2-fold stronger binding of ELK1 in heterozygous variant carriers as compared to reference allele carriers (Fig. 3D). These results demonstrated that the longevity-associated SMAD3 rs8040749 variant enhanced the chromatin binding with ETS family genes such as ELK1, which may lead to altered enhancer activity.

### Longevity-associated SMAD3 haplotype disrupted enhancer activity

To test the impact of longevity-associated SMAD3 regulatory variant on enhancer activity, we performed luciferase reporter assays with reporter constructs harboring SMAD3 rs8040749 major-A or minor-G allele in EBV-transformed human lymphocytes (LCLs). At basal level, no significant allele-specific differences in luciferase reporter activities were detected, although the longevity-associated G-allele showed a trend towards reduced luciferase activity. Furthermore, we treated LCLs with retinoic acid which can upregulate SMAD3 expression in blood cells and found a 2.5-fold increased reporter activity from the major allele construct as compared to the empty vector, whereas the longevity-associated G-allele construct showed significantly decreased reporter activity (P < 0.01) (Fig. 3D). This result suggests that the centenarian-enriched rare SMAD3 variant reduces SMAD3 expression through the disruption of SMAD3 enhancer activity.

### Reduced SMAD3 expression retards cellular senescence in human fibroblasts

To understand the age-related change of SMAD3, we examined SMAD3 expression in 50 human tissues of donors aged from 20 to 70 years from Genotype-Tissue Expression project (GTEx) database^14^. Interestingly, SMAD3 gene expression levels went up significantly with age in lung (R=0.31, P=5e- 10), skeletal muscle (R=0.22, P=1.4e-06), and whole blood (R=0.22 P=2.6e-05) (Fig. 4A). Together with our finding that SMAD3 mRNA level increased dramatically in senescent human fibroblasts (Fig. 4B), these results demonstrate the aging/senescence-associated upregulation of SMAD3. To determine whether reduced SMAD3 expression exerts a beneficial effect to counteract tissue or cellular aging, we knocked-down (KD) SMAD3 in human fibroblasts using shRNA stably delivered by lentiviral transduction and assessed the effect on cellular senescence (Fig. 4C). Upon serial passaging, fibroblasts transduced with a scramble shRNA ceased growth after 99 days, whereas SMAD3-KD fibroblasts maintained a robust proliferation until 135 days (Fig. 4D). In contrast to control fibroblasts, which exhibited progressive senescence, significantly lower percentages of senescence-associated (SA)- β-gal-positive cells (Fig. 4E) and repressed senescence-associated changes in nuclear lamina protein Lamin B1 and cell- cycle regulator p21 ^Cip^^1^ (Fig. 4F) were observed in SMAD3-KD fibroblasts.

**Figure 4.**
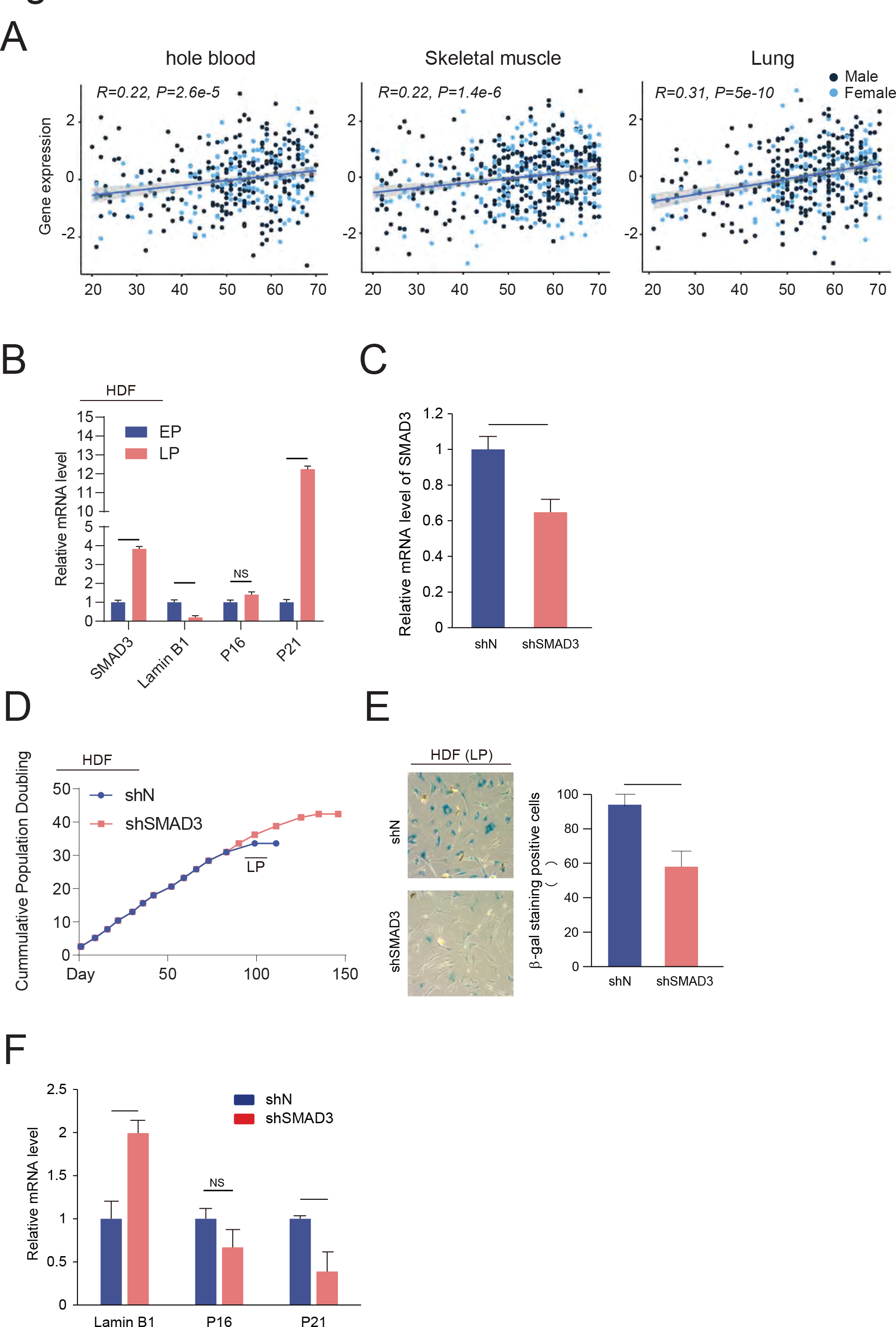
**Identification of rs8040749 as the causal variant** A. Age-dependent changes in SMAD3 gene expression in whole blood, skeletal muscle and lung using GTEx database. B. Cellular senescence-associated SMAD3 changes in human dermal fibroblast (HDF). C. RT-qPCR analysis showing the decreased SMAD3 mRNA level in SMAD3- shRNA transduced HDF. **, p < 0.01. D. Growth curve showing the accumulative population doubling of HDF transduced with non-target or SMAD3 shRNA. E. SA-β-gal staining of HDF transduced with non-target or SMAD3 shRNA at late passage (LP). **, p < 0.01. F. RT-qPCR analysis of senescence-associated markers Lamin B1, P16 and P21. NS, not significant; *. P < 0.05; **, p < 0.01.

### Cell-type specific repression of SMAD3 expression by SMAD3 rs8040749 variant

Our results have demonstrated the longevity-associated rs8040749-G allele created a binding site of ETS family genes, enhanced the binding with ELK1 and decreased enhancer activity in an exogenous enhancer reporter assay. Meanwhile, the downregulation of SMAD3 repressed cellular senescence in human fibroblasts. However, the direct evidence supporting the causal relationship between longevity-associated SMAD3 variants and the transcriptional repression of endogenous SMAD3 expression is still missing. Therefore, we introduced the rs8040749 G-allele into iPSC using CRISPR/Cas9-mediated gene editing to generate a heterozygous line (SMAD3^Cent/+^ iPSCs) (Fig. 5A and S5A). The SMAD3^Cent/+^ iPSCs expressed similar pluripotency makers including OCT4, SOX2 and SSEA4 as compared to it isogenic control (Fig. S5B). The SMAD3 mRNA levels were comparable in SMAD3^Cent/+^ and SMAD3^+/+^ iPSCs (Fig. 5B), which is consistent with the longevity-associated variant-residing enhancer being inactive in PSCs. We further differentiated iPSCs into CD31^+^/CD144^+^ endothelial cells (EC) (Fig. S5C), CD34^+^ hematopoietic stem cells (HSC) (Fig. S5D) and SOX2^+^/Nestin^+^ neural progenitor cells (NPC) (Fig. S5E). SMAD3 mRNA levels were reduced in the gene-edited EC and HSC, but not in NPC (Fig. 5B), consistent with the status of H3K27Ac enhancer activities observed at rs8040749 locus in these cell types. In addition, we performed CRISPRi using dCas9-KRAB in two enhancer-active cell types such as bone-marrow derived MSC and IMR90 fibroblasts. Although these cells didn’t carry the longevity-associated variant, mRNA levels of SMAD3 decreased considerably when the variant-residing region was repressed by dCas9-KRAB (Fig. S5F). Taken together, the cell-type specific repression of SMAD3 caused by longevity-associated variants demonstrated that rs8040749-G allele specifically inhibits an active enhancer. To determine if there are any epigenetic changes caused by rs8040749, we measured allele-specific enhancer activity at the rs8040749 locus in SMAD3^Cent/+^ ECs by digital PCR (Fig. 5C). Fewer DNA fragments harboring the rs8040749-G allele were found in H3K27Ac antibody-pulled down samples, whereas same amount of DNA fragments harboring A allele and G allele in genomic DNA (Fig. 5C). This suggests the SMAD3 longevity-associated variant disrupts the endogenous enhancer at rs8040749 locus. Similarly, we observed fewer DNA fragments harboring rs8040749-G allele than A-allele in ATAC-seq library (Fig. 5C), indicating the chromatin accessibility decreased at rs8040749-G allele comparing with A allele. Using this isogenic stem cell model, our results directly establish the causality of identified SMAD3 regulatory variant rs8040749 in reducing SMAD3 enhancer activity and transcriptional expression thereby contributing to human longevity.

**Figure 5.**
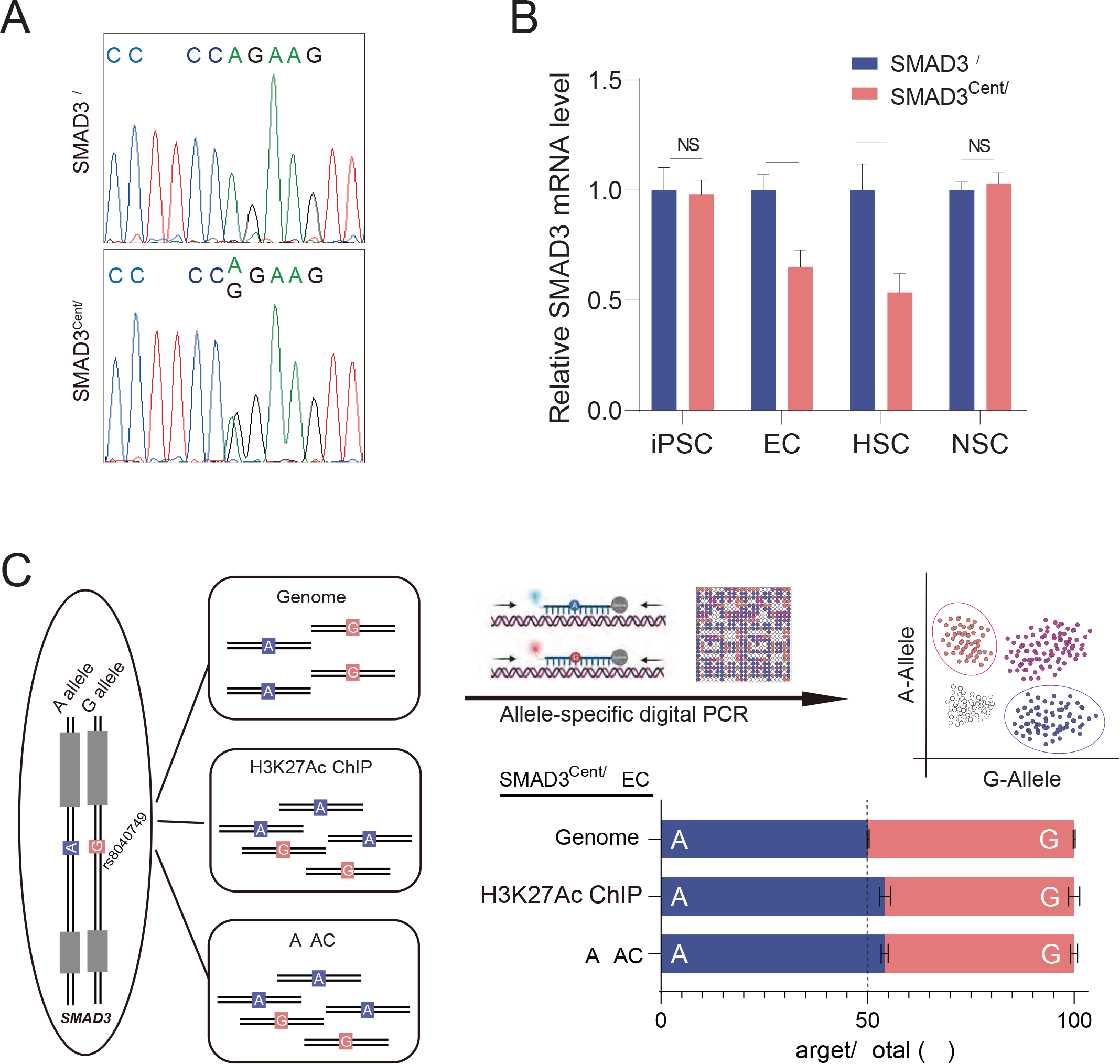
**Identification of rs8040749 as the causal variant** A. Genotypes of SMAD3^+/+^ and SMAD3^Cent/+^ hiPSCs at rs8040749 locus B. RT-qPCR analysis showing the SMAD3 expression in SMAD3^+/+^ and SMAD3^Cent/+^ hiPSCs and iPSC-derived EC, HSC and NPC. NS, not significant; **, P < 0.01; ***, p < 0.001. C. Allele-specific H3K27Ac activity and chromatin accessibility at rs8040749 in SMAD3^Cent/+^ ECs.

### Endothelial cells carrying *SMAD3* rs8040749 variant showed restrained inflammation and resistance to cellular aging

To determine the molecular mechanism by which the SMAD3 variant improved the functionality of vascular cells, we performed transcriptome analysis in EC. Surprisingly, the single nucleotide variant in an enhancer region that reduced SMAD3 expression significantly changed the expression of a large number of differentially expressed genes (DEGs) (FC>2, FDR<0.05) (Fig. S6A). Gene set enrichment analysis revealed downregulation of interferon signaling pathway in SMAD3^Cent/+^ ECs (Fig. 6A and 6B). Interferon regulatory factors and numerous interferon-induced genes were decreased in SMAD3^Cent/+^ ECs compared to SMAD3^+/+^ ECs (Fig. S6B). Consistently, RT-qPCR analysis validated that interferon-induced genes were downregulated in SMAD3^Cent/+^ ECs, but not in SMAD3^Cent/+^ iPSCs when comparing with their isogenic controls (Fig. 6C). These results demonstrate the longevity-associated variant in SMAD3 restrains chronic inflammation in EC.

**Figure 6.**
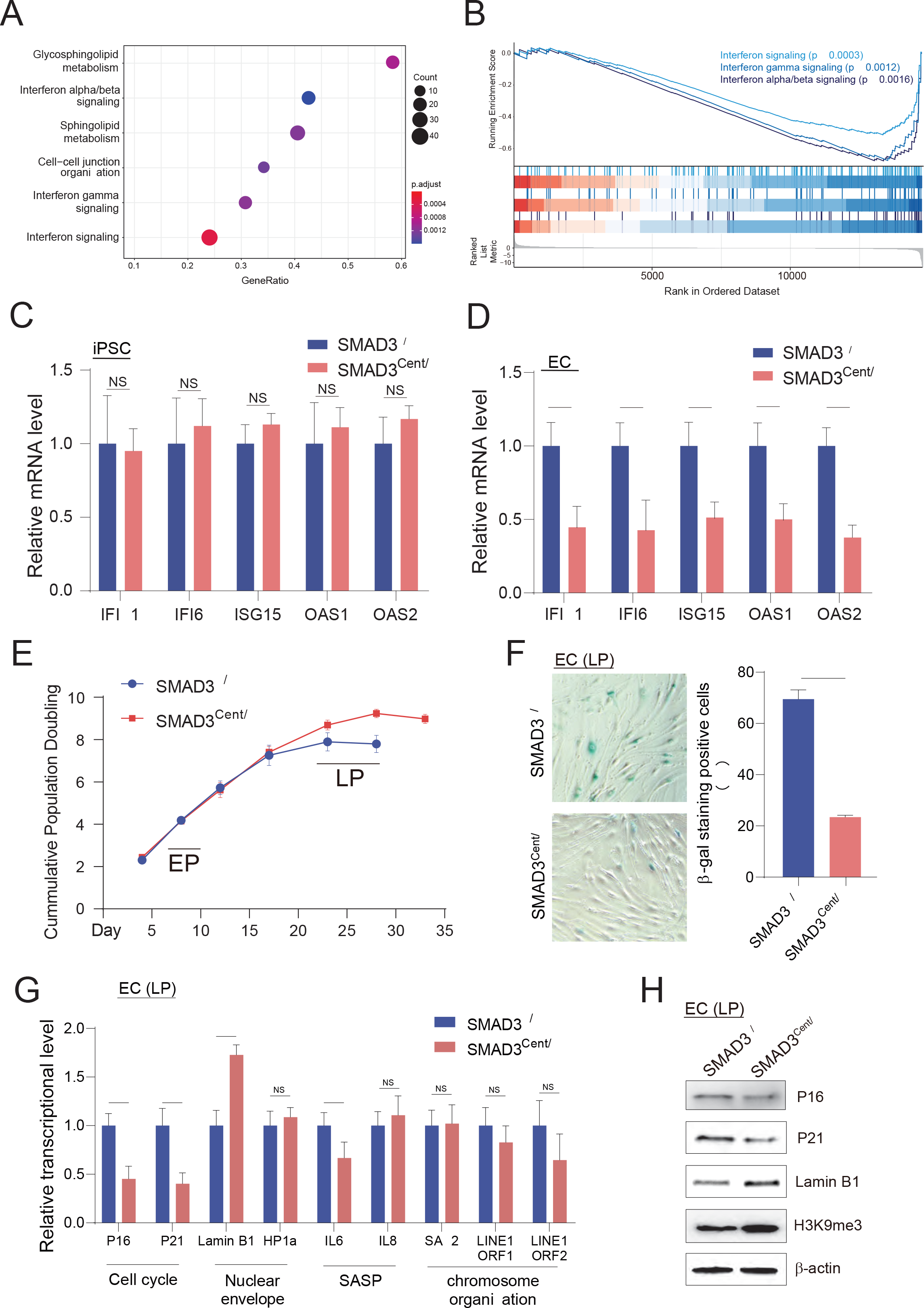
**Functional analysis of SMAD3^+/+^ and SMAD3^Cent/+^ EC** A. Dot plot showing the pathways that were significantly altered by SMAD3 variants in EC. B. The representative downregulated interferon signaling pathways in EC. C. RT-qPCR analysis of interferon signaling genes in iPSC. NS, not significant. D. RT-qPCR analysis of interferon signaling genes in EC. *, P < 0.05; **, p < 0.01. E. Growth curve showing the accumulative population doubling of SMAD3^+/+^ and SMAD3^Cent/+^ EC. ***, p < 0.001. F. SA-β-gal staining of SMAD3^+/+^ and SMAD3^Cent/+^ EC at late passage. ***, p < 0.001. G. RT-qPCR analysis of senescence-associated markers in SMAD3^+/+^ and SMAD3^Cent/+^ EC at late passage. NS, not significant; *, P < 0.05; **, p < 0.01. H. Western blotting analysis of senescence-associated markers in SMAD3^+/+^ and SMAD3^Cent/+^ EC at late passage.

We further investigated the cellular outcomes of the longevity-associated SMAD3 variant in ECs, focusing on the impact on replicative senescence of ECs. Similar to the impact of SMAD3 repression in fibroblasts, SMAD3-edited ECs showed extended *in vitro* lifespan (Fig. 6E) and delayed senescence-associated phenotypes including accumulation of SA-β-gal positive cells (Fig. 6F), upregulation of p16^INK4a^ and p21^Cip^^1^, downregulation of Lamin B1, upregulations of the SASP gene IL6, and loss of heterochromatin marker H3K9me3 (Fig. 6G and 6H) when compared to wild-type ECs.

### Heterozygosity in Smad3 in *Ercc*^1^*^-/^***^Δ^**mice extended healthspan and reduces senescence

To examine the function of longevity-associated SMAD3 variant *in vivo*, we first investigated if the variant-residing region was conserved among different species. The variant-residing region of Smad3 is not conserved with the binding site of ETS-family genes missing in rodents (Fig. 7A). Therefore, we were unable to model directly the function of variant by knocking-in in the mutation or modulating the enhanced binding of ETS-family genes. Instead, we mimicked the directional impact of longevity-associated SMAD3 variant in the Ercc1^−/Δ^ mouse model of accelerated aging that recapitulates human XFE progeria with a median life span of about 26 weeks, by heterozygosity in SMAD3. Health assessments were performed twice per week with tremor, kyphosis, dystonia, ataxia, gait disorder, hindlimb paralysis and forelimb grip strength scored separately. The combined symptom score including signs of aging reflects the overall health condition of Ercc1^−/Δ^ mice. Strikingly, downregulation of SMAD3 significantly extended healthspan especially between 14 and 15 weeks in Ercc1^−/Δ^ mice (Fig. 7B). In addition, reduced p16^INK4a^, p21^Cip^^1^ and SASP genes such as Il-6 and Mcp1 were observed in liver (Fig. 7C), indicating reduced Smad3 expression reduced cellular senescence *in vivo*.

**Figure 7.**
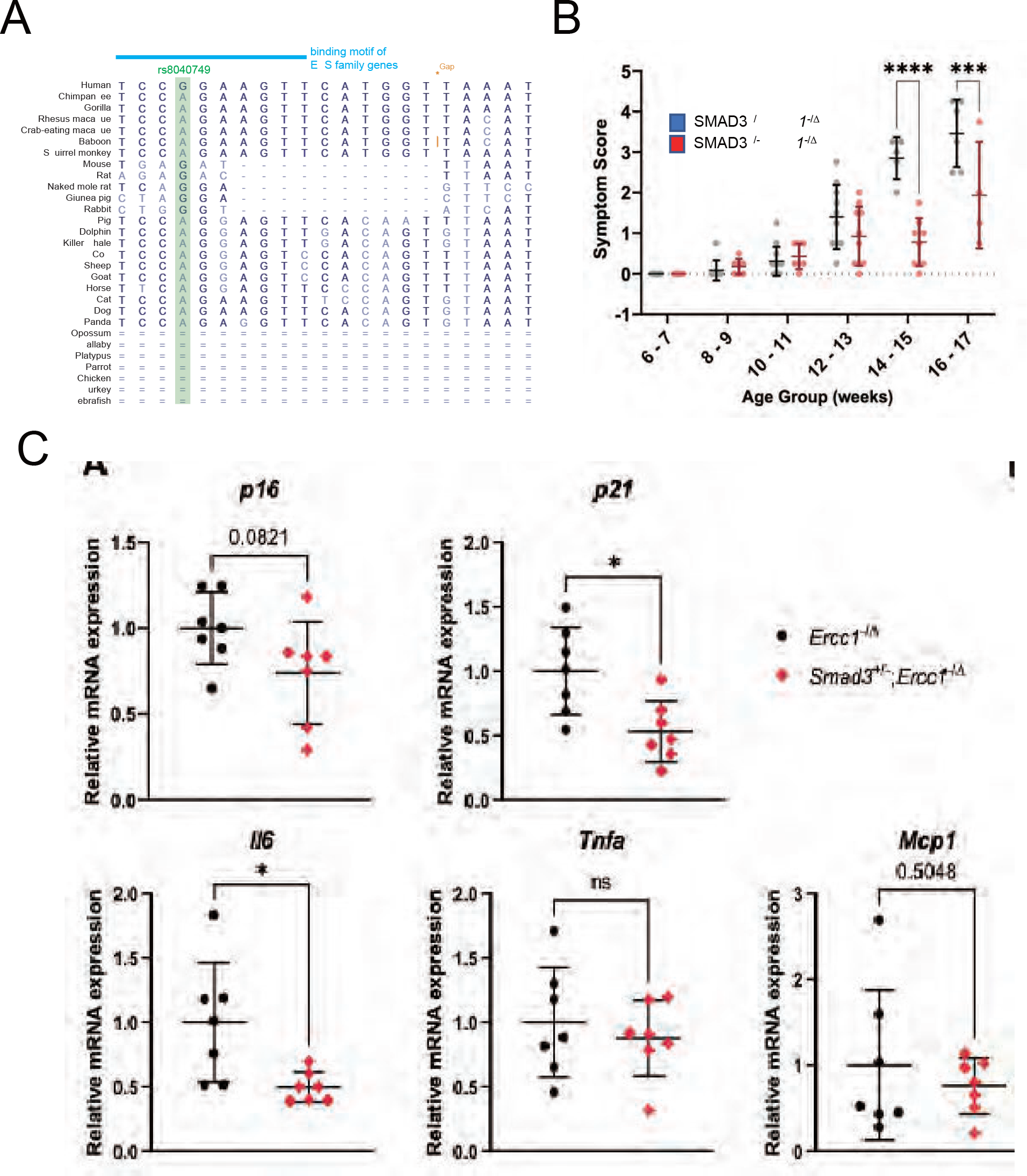
**Heterozygosity in Smad3 in Ercc1-/**Δ **mice extended healthspan and reduces senescence** A. Genomic sequence alignment and conservation among different species at human rs8040749 locus. B. The combined symptom score in Ercc1-/Δ mice. C. RT-qPCR analysis of p176, p21 and SASP factors Il6, Tnfa and Mcp1 in Ercc1-/Δ mice. NS, not significant; *, P < 0.05.

## Discussion

The transforming growth factor-β (TGF-β) superfamily are pleiotropic cytokines that regulates a myriad of cellular processes and recent reports have directly and indirectly linked TGF-β signaling to aging^15, 16^. For example, TGF-β signaling modulates the expression of downstream target genes involving in multiple aspects of aging processes, such as cell cycle regulation, the production of reactive oxygen species (ROS) and inflammatory factors, DNA damage repair, telomere regulation, unfolded protein response (UPR), and autophagy. Previously, heterochronic parabiosis-based aging studies identified certain geronic factors, particularly systemic secretory proteins, that can recapitulate some of the phenotypes observed in heterochronic parabiosis. These identified systemic geronic factors are either members of TGF-β superfamily or are directly modulated by TGF-β signaling pathway^2–5^. Therefore, we hypothesized that human longevity involves genetic variation in TGF-β superfamily signaling genes. Thus, the goal of this study was to identify and characterize functional variants associated with human longevity. We utilized a candidate functional genomic approach and conducted a multi-stage systematic study which included 1) targeted sequencing of 113 genes involved in aging-associated TGF-β signaling in a centenarian cohort; 2) function annotation and prioritization of nominally significant variants within a holistic framework which incorporates genomic and epigenomic information; and 3) comprehensive functional characterization of a longevity-associated novel rare haplotype as well as the causal regulatory variant in SMAD3 to understand the biological mechanisms that underlie genetic association.

Exceptionally long-lived individuals compose only a tiny proportion of the world’s population, but their genomes contain the codes that determine healthy aging and longevity^6, 17^. Previous genetic association studies have yielded only *APOE* and *FOXO3* as significantly correlated with human longevity, likely due to a lack of statistical power^18^. To facilitate the discovery of truly causal variants, we utilized the Ashkenazi Jewish population that has remained genetically isolated for a prolonged period and has restricted gene flow with neighboring populations. The high genetic homogeneity of the isolated population increases the power for association studies for variants, especially rare variants. This population has been used to identify longevity-related genes including functional coding variants in *IGF1R* ^19^ and *SIRT6* ^20^.

Similar to the other genetic association studies, the vast majority (>90%) of variants discovered in genes in the TGF-β pathways in our study reside in non-coding regions of the genome. Given that over 90% of longevity-associated non-coding variants (nominal P < 0.01) were located in *cis*- regulatory elements in at least one cell/tissue type, our candidate sequencing approach showed reasonably good power in the discovery of putative functional variants. Besides, we identified four longevity-associated genes (TGFBR2, SMAD7, TSC22D1 and SMAD3) because of the enrichment of putative regulatory variants in these genes. TGFBR2 encoded one of the three TGF-β receptors in human. Among all variants in TGFBR2, the centenarian-enriched T allele of the common variant (rs4586858) is associated with declined TGFBR2 expression in eQTL, suggesting that repressed TGF-β signaling could be beneficial. Another rare non-coding TGFBR2 variant (rs72850810) is located in the center of a robust enhancer peak. We also found two centenarian-enriched intronic SMAD7 variants (rs141885702 and rs78026276) overlapped with enhancer marks. TSC22D1 can be stimulated by TGF-β is a tumor suppressor crucial for oncogene-induced senescence. We also identified two novel rare centenarian-enriched variants in TSC22D1, including one coding variant in exon 1 (chr 13:45148957) and the other non-coding variant in intron 1 (chr 13:45012396). However, the effect of these variants on function is unknown.

SMAD3, is a transcriptional modulator of TGF-β signaling, acting downstream of TGF-β receptors that is activated by phosphorylation. SMAD3 then binds to SMAD4 and the SMAD3/4 complex translocate to the nucleus to regulate the expression of target genes. Genetic variants in SMAD3 are associated with multiple age-related diseases such as coronary artery diseases, myocardial infarction, osteoarthritis, thyroid cancer, breast cancer, respiratory disease and chronic inflammatory diseases in many human GWAS studies. By both variant-based and function annotation analyses, we identified a significant association of *SMAD3* with longevity. The *SMAD3* locus harbors a large portion of the variants that are enriched in centenarians and the far majority of them are rare in frequency as compared to controls. Furthermore, we found a longevity-associated haplotype in intron 1 of *SMAD3* encompassing 23 centenarian-specific variants that overlapped with a cell type-/tissue-specific enhancer. Surprisingly, the intron 1 of *SMAD3* is quite long (∼ 100 kb) and contains multiple strong enhancers with cell type-/tissue-specificity, suggesting that there are sophisticated chromatin interactions within this region controlling the expression of SMAD3. The novel SMAD3 enhancer we identified was silent in pluripotent stem cells and brain-related cell types, but was active in blood cells, mesoderm cells and fibroblasts. Interestingly, the enhancer within intron 2 of SAMD3 contained GWAS variants associated with asthma, thyroid cancer and coronary artery diseases (CAD) while the intron 4 enhancer contained multiple asthma-associated GWAS variants, both of the enhancers showed different enhancer activities. In addition, we found the H3K27Ac peaks showed different patterns between blood cells and other cells, probably due to a unique binding profile of TFs and co-factors in blood cells.

The longevity-associated G allele of rs8040749 from the haplotype created a binding site for ETS family genes, whereas the reference A allele disrupted it. The ETS family genes are transcriptional factors and share the same consensus binding motif. Based on the TF ChIP-seq from ENCODE, both ELF1 and ELK1 bind to the motif at SMAD3 rs8040749 locus in lymphoblastoid cell line GM12878. Thus it is highly likely that multiple ETS genes can competitively bind to the motif in the novel SMAD3 enhancer. Moreover, it is known that ELK1 can act as a transcriptional activator or repressor^21^, depending on its binding with a coactivator or corepressor, while other ETS family genes may have either single or dual role in transcriptional regulation. Hence knowing the variant-associated TF binding change is not sufficient to predict the directional impact of the longevity-associated variants. Our enhancer reporter assays indicated that the directional impact of the longevity-associated variant was to reduce SMAD3 expression, which was the opposite direction of aging- and senescence- associated changes. Consistently, GWAS variants associated with lower risk of CAD were associated with lower SMAD3 levels^22^. Mouse studies also showed diminished adiposity with improved glucose tolerance and insulin sensitivity ^23^ as well as resistance to fibrosis across several tissues^24–27^ in SMAD3 null mice. Since complete knock-out of SMAD3 led to death at 8 months in mice, we believe the presence of SMAD3 is important to maintain essential cellular process such as immune function, but reduced SMAD3 could exert beneficial effect in certain tissues in aging and disease contexts.

Due to the lack of studies on biological mechanisms that underlie genetic association, the genetic basis of human longevity is still poorly understood. So far, only limited mechanistic studies of longevity-associated variants have been reported. For example, a BPIFB4 coding variant enriched in long-living individuals reduced endothelial dysfunction and slowed atherogenic plaque progression in atherosclerosis-prone mice, enhanced vascular function in human endothelial cells and rejuvenated immune system and frailty in aged mice^28–30^. In addition, longevity-associated SIRT6 coding variants showed enhanced mono-ribosylation activity, improved DNA repair function and cancer killing capability in human cells^20^. However, since the non-coding variants are in regions that lack sequence conservation among species, elucidating functionality of regulatory variants has proven to be even challenging than coding variants. Given the intricate nature of genome organization, which involves DNA, histones, transcription factors, noncoding RNAs, and long-range chromatin interactions, the commonly employed reporter assay may not accurately reflect the true effects of regulatory variants^31^. Therefore, we examined function of the non-coding variants in an endogenous genome in variant edited iPSCs and in lineage-committed differentiate cell types. The modeling of longevity using PSCs and their differentiated cell types in our study demonstrated the cell-type specific repression of SMAD3 which was in line with the enhancer activity in different cell types. In addition, we were able to unbiasedly detect allele-specific enhancer activity and chromatin accessibility using the heterozygous cells. The variant-mediated lowered SMAD3 enhancer activity repressed SMAD3 expression and further led to the dampened interferon signaling. It has been reported that SMAD3 interacts with Interferon Regulatory Factor 7(IRF-7) to activate IFN transcription^32^. Additionally, upregulation of genes in sphingolipid metabolism and cell-cell junction organization were observed in the SMAD3- edited ECs. Lastly, we demonstrated the rare variant reduced replicative senescence in ECs.

Since the non-coding region where rs8040749 resides is conserved in primates, but not in rodents, it isn’t feasible to model the variants in mouse models. Instead, we mimicked the effect of SMAD3 variants in the Ercc1^-/Δ^ progeroid mouse model of accelerated aging^33^ by generating mice heterozygous for SMAD3. Here reducing SMAD3 levels in all cell types and tissues results in an extension of healthspan and a reduction in senescence and SASP expression in multiple tissues. Taken together, our cell culture and *in vivo* results are consistent with a reduction in SMAD3 expression conferring a reduction in senescence, SASP and other markers of inflammation. Thus development of approaches to reduce SMAD3 levels such as through the use of small molecules targeting SMAD3 or phosphorylation of SMAD3 or the application of CRISPRi-based approaches to suppress SMAD3 expression should extend human healthspan.

## Materials and Methods

### Study subjects and sample collection

This study group consisted of 1,068 Ashkenazi Jewish samples: 496 Ashkenazi Jewish centenarians and 572 Ashkenazi Jewish controls that were previously collected as part of longevity study at the Albert Einstein College of Medicine^1^. A centenarian is defined as a healthy individual living independently at 95 years of age or older and a control is defined as an individual without a family history of unusual longevity; parents of controls survived to the age of 85 years or less. Informed written consent was obtained in accordance with the policy of the Committee on Clinical Investigations of the Albert Einstein College of Medicine. Genomic DNAs were extracted from blood samples and amplified using illustra GenomiPhi V2 DNA Amplification kits (GE healthcare Life Sciences).

### Selection of target genes and generation of customized target capture

113 candidate genes implicated in TGF-β signaling includes (1) TGF-β superfamily members such as TGFBs, Growth and Differentiation Factors (GDFs), Bone Morphogenetic Proteins (BMPs), Myostatin, Activins/Inhibins and Nodals; (2) TGF regulators such as ligands, antagonists and inhibitors; (3) signal receptors and co-receptors; (4) signal transducer SMADs, SMAD regulators, transcriptional factors and transcriptional targets. The information of candidate genes was shown in Table S1.

For 23 of the selected genes, we included the whole gene and 2 kb upstream from the transcription start site (TSS), except for GDF11, where the region 10 kb upstream from the TSS was included. For the rest genes, we included the 2 kb region upstream of the TSS, all exons, and 20 bp of each exon-intron junction. We designed a customized Nimblegen SeqCap EZ Choice library (Roche) for target capture of the 113 gene regions mentioned above using the Nimbledesign tool (https://design.nimblegen.com/nimbledesign) with ‘Preferred close match’ set to 3 and ‘Maximum close match’ set to 20. The targeted regions totaled 2,199,107bp with 95.3% base coverage.

### Capture sequencing of TGF-**β** signaling-associated genes

The sequencing of these 113 genes in AJ centenarian cohort was similar as previously described^1–3^.

For the 20 pools of centenarians and 23 pools of controls, 10/11-plex pre-pooled target capture was performed and sequenced in 43 lanes by Illumina HiSeq 2000. Libraries were prepared using the TruSeq DNA sample preparation version 2 low-throughput (LT) protocol from Illumina with the following modifications: 1) A Covaris system was used to shear DNA to 300bp

1. Each Sample was ligated using 1 of the 12 unique Illumina Truseq adapters.
2. A 0.8X volume of AMPure XP beads (Beckman Coulter) were used in two steps for the initial clean-up. 4) Size selection was performed by using a 0.65X volume of AMPure XP beads and collecting the supernatant. It was mixed with a 0.85X volume of the beads, which were washed by 70% ethanol and suspended in resuspension buffer. Pre-capture enrichment of the eluted DNA from the beads was performed by PCR amplification for 7 cycles and PCR products were purified using 0.85X volume of AMPure XP beads.

Four target capture reactions were performed (3X11 plex and 1X10 plex) using the Nimblegen SeqCap EZ Choice library customized for our candidate genes as per the Agilent SureSelect Target Enrichment protocol. Equimolar amounts of 10/11 post-enrichment libraries were pooled to 1ug. As per the modified protocol, six index blocking oligos from IDT were resuspended in 300uM water and further diluted to 50uM using the Indexed Blocking Reagent (IRB). Post-capture PCR was performed for 12 cycles and libraries were sequenced by Illumina HiSeq 2000 (paired-end 100bp).

### Variant-based analysis

The sequencing reads were aligned to the human reference genome assembly (GRCh37/hg19) using BWA^4^. We performed quality control to remove low-coverage reads and PCR duplicates prior to the alignment. The Picard tools CollectAlignmentSummaryMetrics and CollectTargetedPcrMetrics, version 1.81, were used to calculate the read coverage in each pool. Variants were called using the software program CRISP, specifically developed to call common and rare variants in pooled DNA sequence data^5^. Genetic variants were annotated with dbSNP (version 150) using ANNOVAR^6^.

### linkage disequilibrium (LD) analysis

Linkage disequilibrium (LD) heatmap at *SMAD3* locus was generated from 1000 Genomes Project VCF files by LDBlockShow 1.40 ^7^. The linked variants were identified in European population by LDproxy Tool (https://analysistools.cancer.gov/LDlink/?tab=ldproxy).

### Luciferase reporter assays

LCLs were treated with 2nM Retinoic acid for 24 hours prior to transfection of firefly and renilla luciferase constructs by Amaxa Nucleofector kit V (Lonza) with the X001 program on a Nucleofector II device. The firefly luciferase constructs were co-transfected with the Renilla luciferase vector (Promega) in 1-2X10^6 LCLs at a ratio of 2ug:40ng respectively. 36 hours after transfection firefly and renilla luciferase activities were measured using the Dual-Luciferase Reporter Assay System (Promega) according to the manufacturer’s protocol, using un-transfected cells to adjust for background activity. Luminescence was measured by Synergy 4 multimode luminometer (BioTek).

### Electrophoretic mobility shift assay (EMSA)

Primers for rs8040749 were designed based on the genomic sequence surrounding the SNP: rs8040749_A-F, 5’-CCTGCCTTCCAGAAGTTCATG-3’, rs8040749_A-R, 5’-CATGAACTTCTGGAAGGCAGG-3’, rs8040749_G-F, 5’- CCTGCCTTCCGGAAGTTCATG-3’, rs8040749_G-R, 5’-CATGAACTTCCGGAAGGCAGG3’. The variable nucleotide is shown in bold. All the primers were ordered from Integrated DNA Technologies with biotinylated 3’ ends. Corresponding forward and reverse primers were mixed in a 1:1 molar ratio and annealed using a thermocycler to create double-stranded probes. EMSA reactions were performed with the Lightshift chemiluminescent kit (Thermo fisher scientific, 20148) according to the user manual, with 8ug of LCL nuclear extract prepared using the NE-PER Nuclear and cytoplasmic extraction kit (Thermo Fisher Scientific, 78833) and 30 ng of recombinant purified ELK1 (Sino Biologicals 12621-H20B). For competition assays, we used a 200-fold excess of unlabeled probe. The protein complexes were resolved on 6% DNA retardation gels (Invitrogen) for 1 h at 100 V, transferred to Biodyne B Nylon Membranes (Pierce) for 45 minutes at 100 V, crosslinked, and processed with the Chemiluminescent Nucleic Acid Detection Module (Pierce).

### Gene editing at *SMAD3* locus

CRISPR/Cas9-mediated knock-in was performed using Alt-R CRISPR-Cas9 System (IDT) and HDR Donor Oligos. Cas9 nuclease, sgRNA (TTTAACCATGAACTTCTGGA) targeting variant site and single-strand DNA oligo donors (ssODNs) containing SMAD3 rs8040749-G were ordered from IDT. To generate SMAD3 knock-in iPSCs, 2×10^5 individualized iPSCs were resuspended in 10 μL Resuspension Buffer R (Invitrogen) containing CRISPR ribonucleoproteins (Cas9 nuclease +sgRNA) and ssODNs, the cell suspension was then electroporated using NEON Transfection System (Invitrogen). After electroporation, cells were seeded on Matrigel-coated plates in mTeSR Plus with 1x RevitaCell Supplement (Gibco). After 48h expansion, cells were dissociated by Accutase and 10,000 cells were seeded on CytoSort™ Array (10,000 microwells, CELL Microsystem). Once cells were attached, microwells containing single colony were automatically picked and transferred to 96-well plate by CellRaft AIR System (CELL Microsystem). The expanded clones on 96-well plate were further genotyped by TaqMan genotyping assay (rs8040749, ThermoFisher) and Sanger sequencing.

### Antibodies

Antibodies for western blotting and immunostaining were purchased from the following companies.

BD bioscience: SSEA4(560126), CD34(555821), CD31(555445), CD144 (560410), p16(550834)

Santa Cruz Biotechnology: OCT4 (sc-5279), β-actin (sc-47778) Millipore: SOX2 (2003601), NESTIN (MAB5326)

Cell Signaling Technology: p21 (2947)

Abcam: Lamin B1(ab16048), H3K9me3 (ab8898)

### Cell culture

Human BJ iPSCs (BJ fibroblasts were originated from ATCC) were maintained on Matrigel (BD Biosciences) in mTeSR Plus medium (STEMCELL Technology).

Fibroblasts (BJ or IMR90) were culture in DMEM (Gibco) with 10% fetal bovine serum (FBS, GeminiBio) and 1% penicillin/streptomycin (Gibco).

Endothelial cells were cultured in EGM2 endothelial cell growth medium (Lonza CC-3162).

Bone-marrow-derived mesenchymal stem cells were cultured in MEMa (Gibco) with 10% fetal bovine serum (FBS, GeminiBio), 1% penicillin/streptomycin (Gibco) and 1ng/uL bFGF.

### Endothelial cell differentiation

Differentiation of iPSCs into ECs was performed as previously described with minor modifications ^8^. Briefly, iPSCs were cultured in mTeSR1-Plus media for one day and then in M1 medium, containing IWP2 (3 mM), BMP4 (25 ng/ml), CHIR99021 (3 mM) and bFGF (4 ng/ml), for three days. The following day, M1 medium was removed and replaced with M2 medium with the addition of VEGF (50 ng/ml), bFGF (20 ng/ml) and IL6 (10 ng/ml) to promote endothelial cell emergence for another three days. The differentiated adherent cells were harvested using TrypLE (GIBCO), labeled with CD144 (VE-Cadherin) MicroBeads (Miltenyi Biotec 130-097-857), and separated by OctoMACS™ Separator.

### Hematopoietic stem cell differentiation

Differentiation of iPSCs into ECs was performed using STEMdiff Hematopoietic Kit (StemCell Technology, 05310). Briefly, dissociated small iPSC aggregates were seeded to Matrigel-coated plate and cultured in mTeSR-Plus media on day 0. At day 1, attached sparse colonies were cultured in Medium A, consisting of Hematopoietic Basal Medium and 1x Hematopoietic Supplement A, for three days (half-medium change at day 3). At Day 4, Medium A was removed and replaced with Medium B, consisting of Hematopoietic Basal Medium and 1x Hematopoietic Supplement B. Half-Medium B change was performed on day6, 8 and 11. On day 13, HSCs were harvest from the culture supernatant and were purified by MACS using CD34 MicroBeads (Miltenyi Biotec, 130-046-702).

### Neural progenitor cells differentiation

Differentiation of iPSCs into NPCs was performed using PSC Neural Induction Medium (Thermo Fisher) according to manual.

### CRISPRi

dCas9-KRAB (#110820) and pCAGmCherry-gRNA (#87110) plasmids were purchased from Addgene. Scramble-targeted sequence (GCTTAGTTACGCGTGGACGA) and SMAD3 rs8040749-targeted sequence (TTTAACCATGAACTTCTGGA) were cloned into pCAGmCherry-gRNA plasmids separately. For bone marrow-derived MSCs, dCas9-KRAB and gRNA plasmids targeting either scramble sequence or SMAD3 rs8040749 were transfected using lipofectamine 3000 (Thermo Fisher). For IMR90 fibroblasts, plasmids were electroporated using Neon transfection system

(1100 V, 30 ms, 1 pulse) (Thermo Fisher). Cells were cultured for 48h and harvested for RT-qPCR analysis.

### ATAC-seq library preparation

Omni-ATAC-seq protocol was used as previously described^9^. Briefly, 50,000 cells were collected and resuspended in 50 μl cold ATAC-Resuspension buffer (10 mM Tris-HCl pH 7.4, 10 mM NaCl, 3mM MgCl2, 0.1% NP40, 0.1% Tween-20, 0.01% Digitonin) for 3 min on ice, nuclei were centrifuged at 500 RCF for 10 min at 4 °C. The cell pellet was resuspended with 50 μl transposition mix containing 25 μl 2X TD buffer, 2.5 μl Tn5 transposase, 16.5ul PBS, 0.5ul 1% digitonin, 0.5ul 10% Tween-20, and 5 μl water. The mixture was incubated at 37 °C for 30 min in a thermomixer with 1000RPM mixing. Transposed genomic DNA was cleaned up using DNA Clean & Concentrator-5 (Zymo Research). The library was amplified using NEBNext high-fidelity PCR master mix containing 1.25 μM customized Nextera universal (Ad1_noMX) and indexed primer with the cycling condition of 98 °C for 30 s, 8 cycles of 98 °C for 10 s, 63 °C for 30 s, 72 °C for 1 min. The amplified library was purified using AMPure XP beads (Beckman). A double size selection was performed with 0.5x/1.8x bead volume to remove amplicons > 1000 bp or < 100 bp. Libraries were subjected to digital PCR to evaluate allele-specific chromatin accessibility at SMAD3 rs8040749 loci.

### Chromatin immunoprecipitation

For chromatin immunoprecipitation, SimpleChIP Enzymatic Chromatin IP Kit (Magnetic Beads) (CST) was used according to manual. Briefly, 10^6 endothelial cells were cross-linked in 1% vol/vol formaldehyde/PBS for 8 min at room temperature and then quenched by Glycine. Samples were lysed on ice for 5 min. Subsequently, lysates were incubated with MNase to digest DNA. The collected supernatants were incubated overnight with Protein A dynabeads (Life technology, 10001D) associated with H3K27Ac antibody (Active Motif). Next the input sample and chromatin-beads complexes sample were digested, eluted and cross-link-reversed at 68 °C for 2 h on a thermomixer. DNA was finally purified using Spin Columns. The enriched DNA was subjected to digital to evaluate allele-specific H3K27Ac modification at SMAD3 rs8040749 loci.

### Growth curve assay

Cell population doubling was determined as previously described. Briefly, human diploid cells including IMR90 fibroblasts and iPSC-derived endothelial cells were serially passaged and the number of cells was counted. Population doubling per passage was calculated as log2 (number of cells obtained/number of cells plated). Cumulative population doublings of the cells were calculated and plotted to days.

### SA-**β**-gal staining assay

The SA-β-gal staining of hMSCs was conducted as previously described^10^.

### Western blot

Cells were lysed in RIPA buffer (Thermo Fisher) with protease inhibitor cocktail (Roche). Protein quantification was performed using a BCA Kit (Thermo Fisher). Protein lysate was subjected to SDS-PAGE and subsequently electrotransferred to a PVDF membrane (Bio-rad). Then primary antibodies and HRP conjugated secondary antibodies were incubated with the 5% milk blocked membrane. The imaging and quantification of target proteins was obtained by ChemiDoc MP imaging system (Bio-Rad).

### RT-qPCR

Total RNA was extracted using RNeasy Mini Plus Kit (Qiagen). Then One-step PrimeScript RT-PCR kit (Takara) was used to generate cDNA. RT-qPCR was performed with PowerUp SYBR Green Master Mix (Thermo Fisher) in QuantStudio 6 Pro Real-Time PCR System (Thermo Fisher). All primer sequences for PCR are given in Table S4.

### Digital PCR

Digital PCR reaction consisting 1x QuantStudio 3D Digital PCR Master Mix (Thermo Fisher), 1x TaqMan Genotyping assay (Thermo Fisher) and templates (genomic DNA, ChIP DNA or ATAC-seq library) were loaded onto QuantStudio 3D Digital PCR 20K Chip and run in ProFlex 2x Flat PCR System (Thermo Fisher). The results were analyzed by QuantStudio 3D AnalysisSuite Cloud Software (Thermo Fisher).

### Immunofluorescence staining

Cells were fixed in 4 % paraformaldehyde at room temperature (RT) for 15 min, permeabilized in 0.4% Triton X-100/PBS at RT for 10 min. After blocking with 10% donkey serum (Jackson ImmunoResearch Labs)/PBS for 1 h, cells and sections were incubated with primary antibodies at 4 °C overnight and the corresponding secondary antibody (Invitrogen) at RT for 45 min. Nuclei were stained with Hoechst 33342 (Thermo Fisher 62249).

### RNA-seq

Total RNA was isolated with the RNeasy Mini Plus Kit (Qiagen) from 10^6 cells per duplicate following the manuals. Then the quality of RNA was checked with Bioanalyzer (Agilent), following by RNA library construction by Ribo-Zero Plus rRNA Depletion Kit (Illumina) according to the manufacture’s protocol. Paired-end reads were generated from Illumina NovaSeq 6000 Sequencing System. The reads were aligned to the human reference (hg38) by STAR 2.7.6a^11^ and counted by featureCounts from Subread 2.0.2 package^12^ at gene level. Differential gene expression was analyzed by

DSEeq2^13^ and assessed for Gene set enrichment analysis by clusterProfiler 4.0^14^.

### Statistical analysis

The statistical analyses were performed using PRISM software (Graphpad Software). Comparisons were performed with two-tail student’s t-test unless otherwise stated. P<0.05 was defined as statistically significant.

### Data availability

All sequencing data have been deposited to Sequence Read Archive in the National Center for Biotechnology Information. The accession numbers were as follows: targeted genome sequencing data (PRJNA669034), endothelial cell RNA-seq data (PRJNA904291).

---

**Figure S1.**
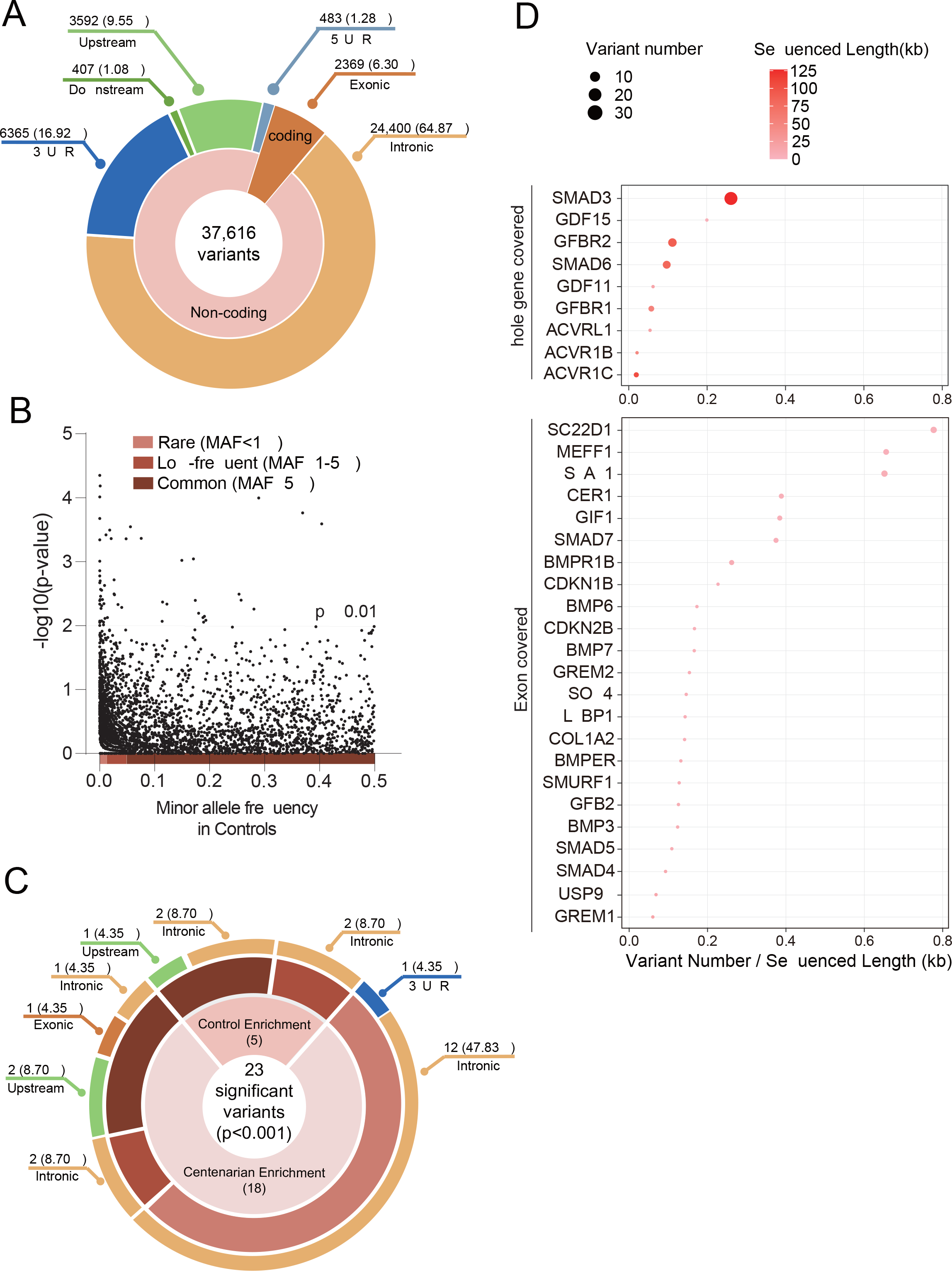
Capture-seq in an Ashkenazi Jewish centenarian cohort A. All 37,616 variants categorized by region. B. Scatter plot showing rare variants tend to show higher significance compared to common variants C. 23 significant variants (P<0.001) categorized by allele frequency and region. D. Dot plot showing the variant discovery rate of each gene. Variant discovery rate means significant variant number per kilobase sequenced length.

**Figure S2.**
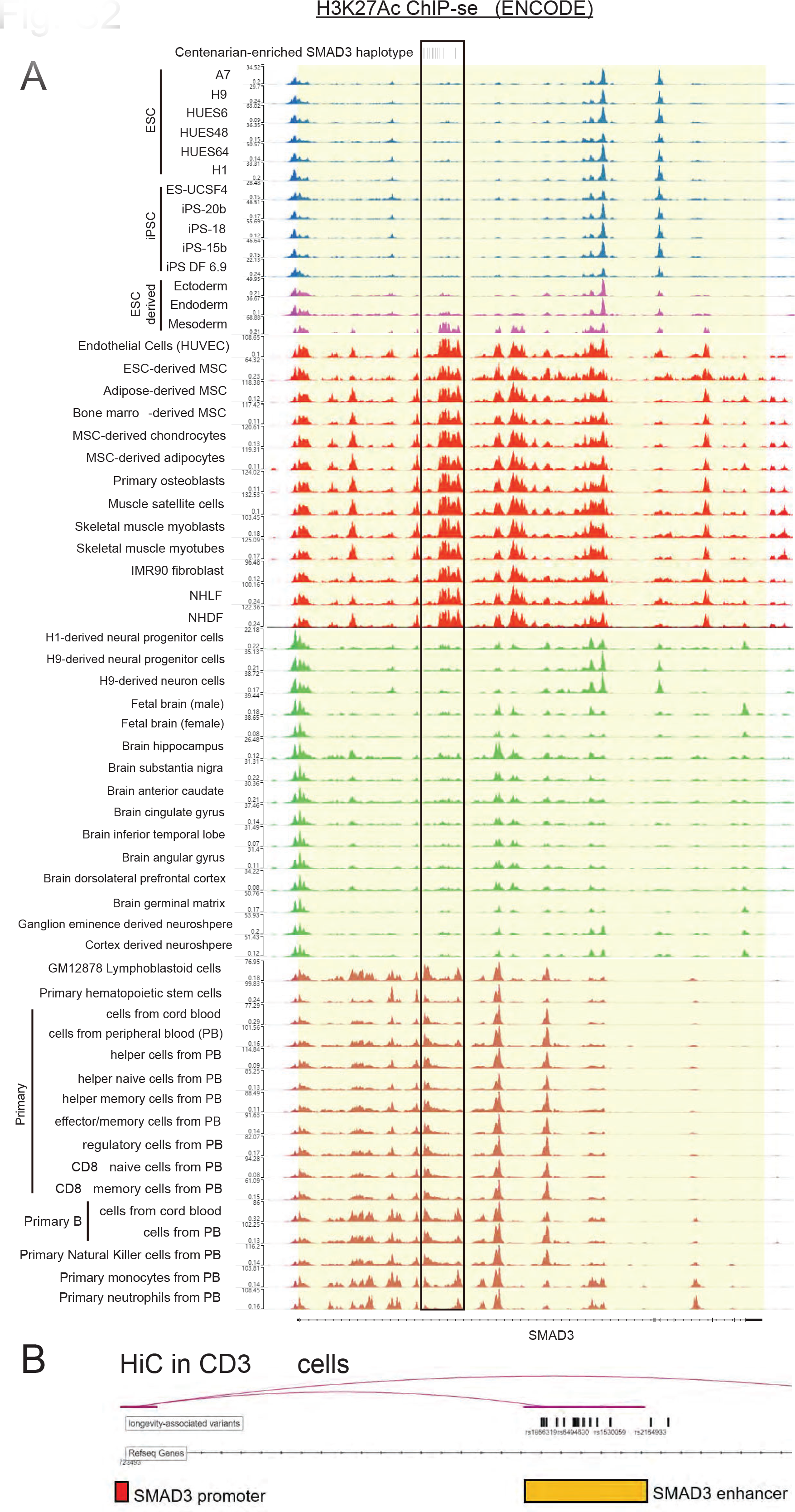
**The longevity-associated haplotype lies in an enhancer looping with SMAD3 promoter** A. Enhancer signals at SMAD3 gene in different cell types/tissues indicated by H3K27Ac ChIP-seq in ENCODE. B. Hi-C data showing the haplotype-residing enhancer in SMAD3 intron 1 directly interacted with SMAD3 promoter in CD3+ T cells.

**Figure S3.**
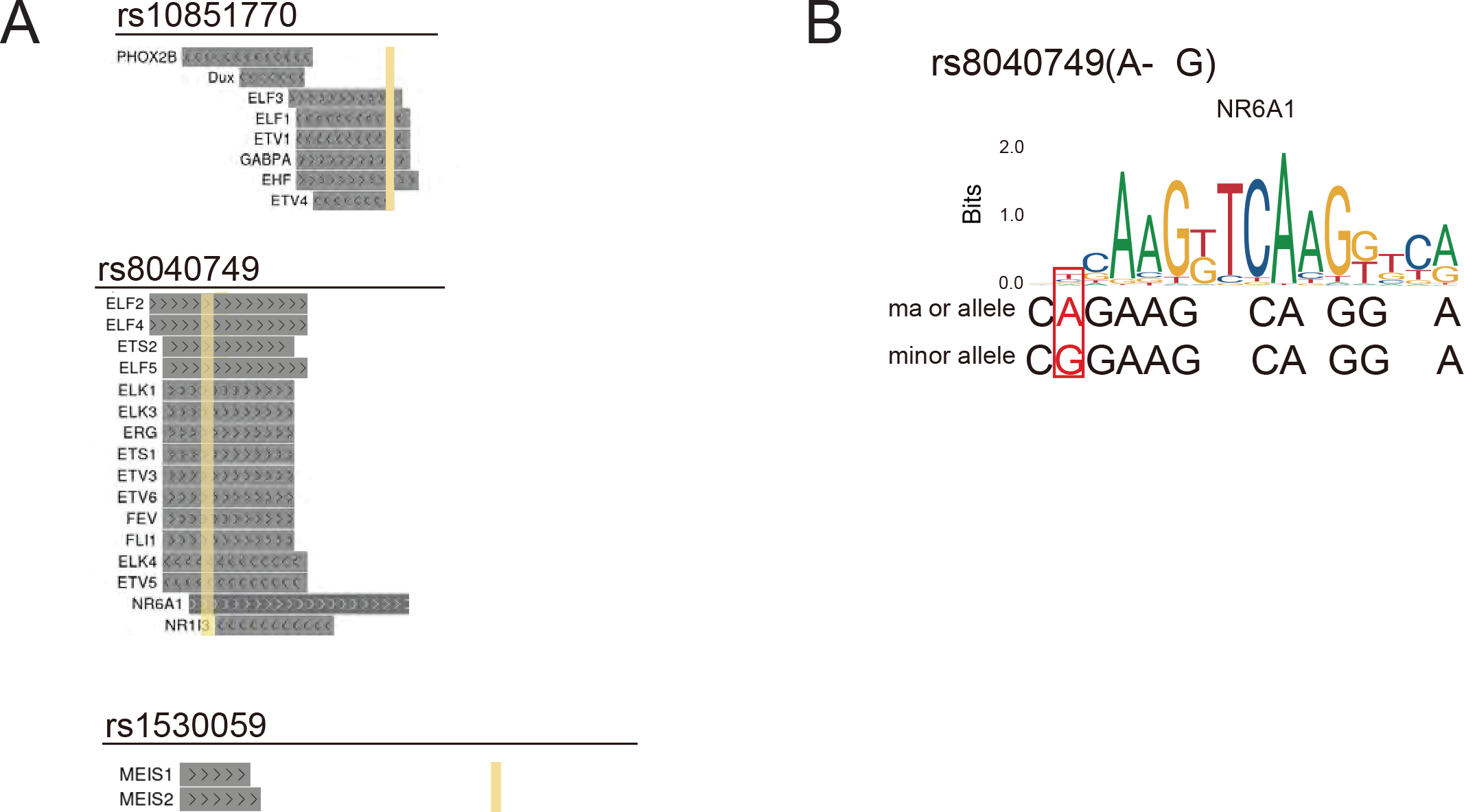
**TF bind prediction of three variants showing Regulome DB score = 2b.** A. The predicted TF binding at rs10851770, rs8040749 and rs1530059 loci. B. rs8040749 overlapped with the binding motif of NR6A1.

**Figure S5.**
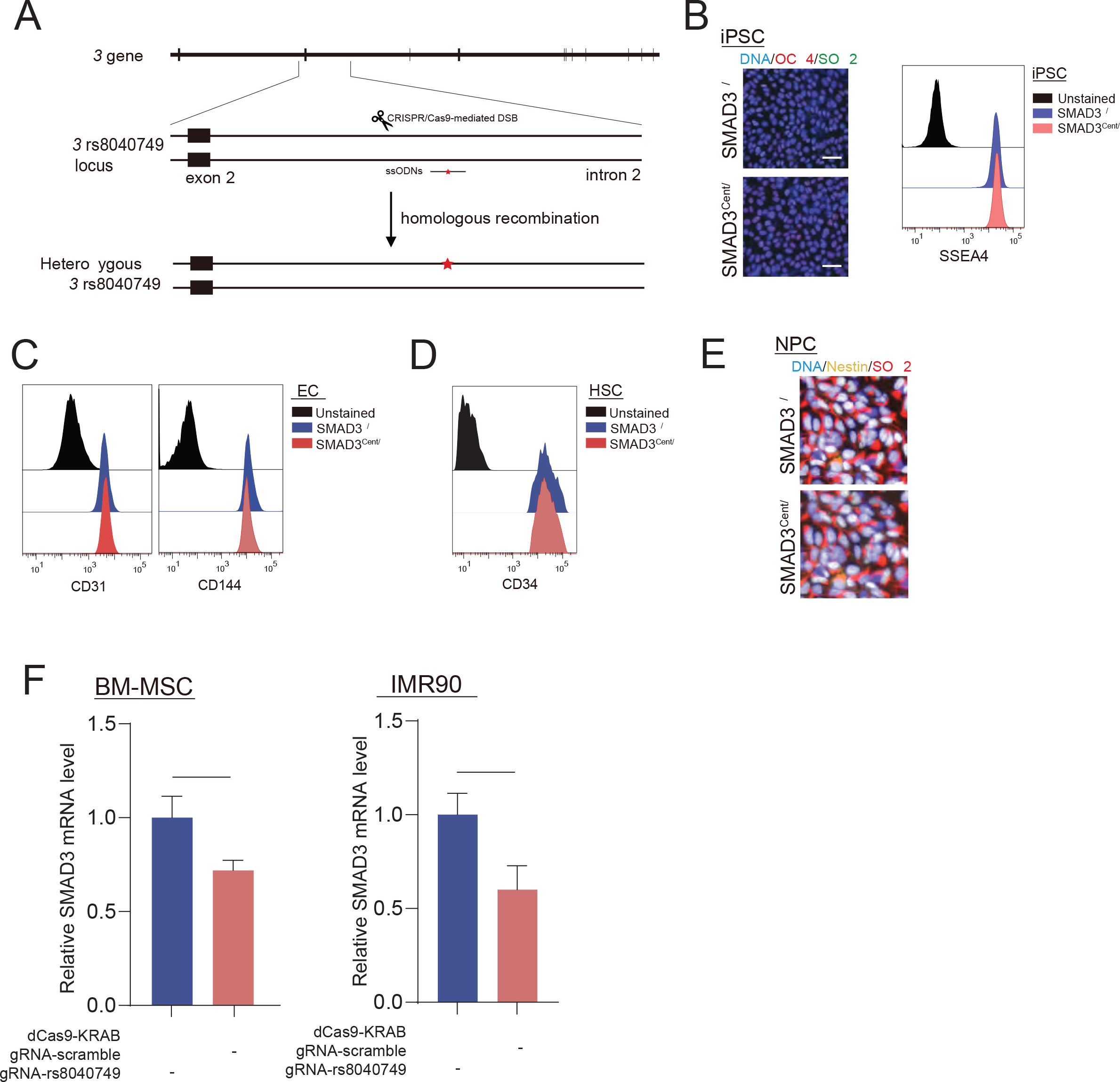
**Characterization of SMAD3^+/+^ and SMAD3^Cent/+^ iPSCs and their derivates.** A. Schematics showing the strategy to knock-in rs8040739-G allele in iPSC. B. Characterization of SMAD3^+/+^ and SMAD3^Cent/+^ iPSC by the detection of OCT4, SOX2 and SSEA4 expression. C. Characterization of SMAD3^+/+^ and SMAD3^Cent/+^ EC by the detection of CD31 and CD144 expression. D. Characterization of SMAD3^+/+^ and SMAD3^Cent/+^ HSC by the detection of CD34 expression. E. Characterization of SMAD3^+/+^ and SMAD3^Cent/+^ NPC by the detection of SOX2 and NESTIN expression. F. RT-qPCR analysis of SMAD3 mRNA level in cells transfected with dCas9-KRAB and scramble gRNA or gRNA targeting rs8040749 locus. *, p<0.05.

**Figure S6.**
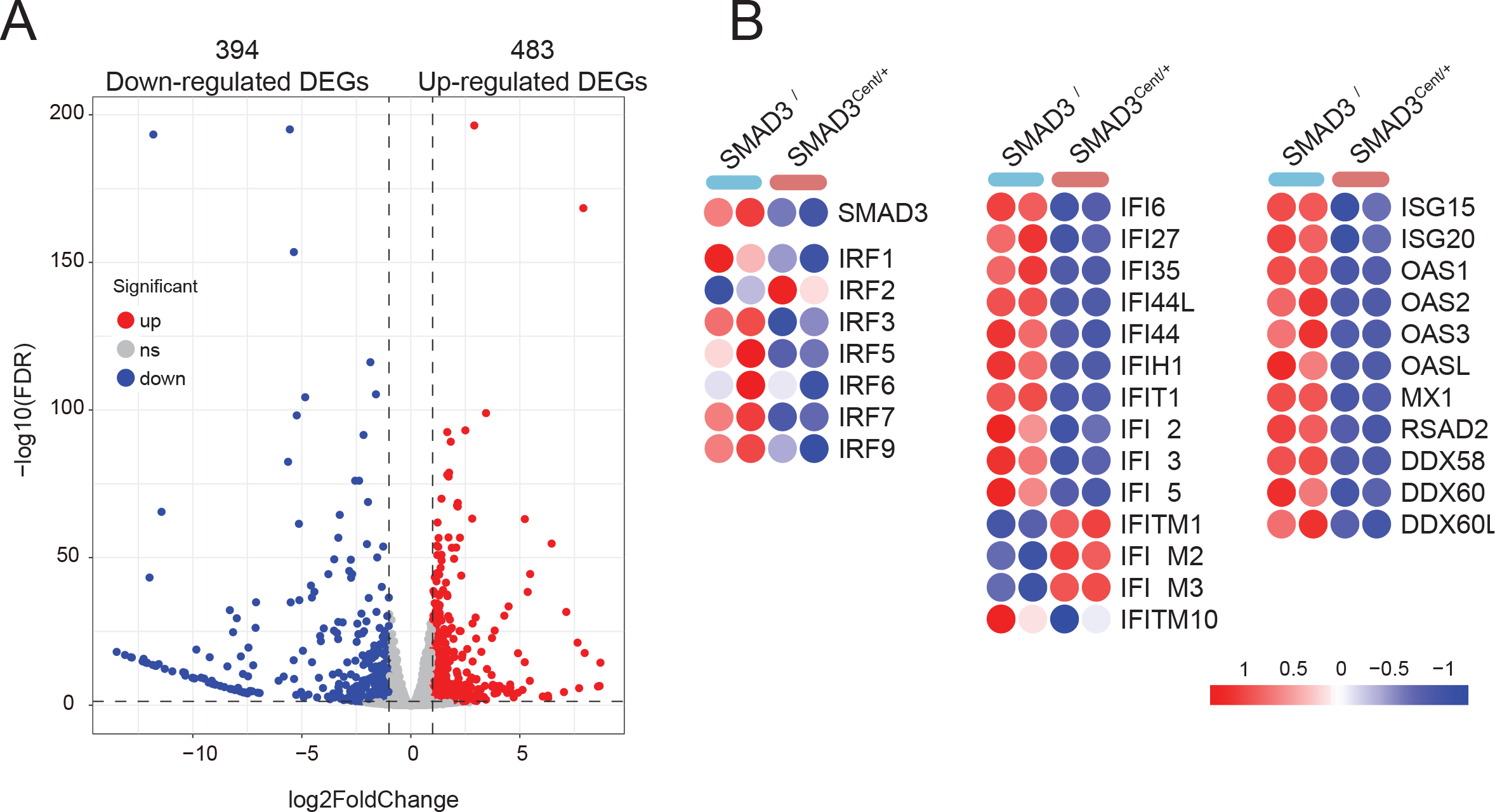
**Transcriptome analysis of SMAD3^+/+^ and SMAD3^Cent/+^ EC** A. Volcano plot showing differentially expressed genes comparing SMAD3^Cent/+^ EC to SMAD3^+/+^ EC. B. Heatmap showing the expression of SMAD3, transcriptional factor IRFs and interferon-stimulating genes in SMAD3+/+ and SMAD3Cent/+ EC.

## Supplementary Tables

Table S1. 113 candidate TGF-β signaling-related gene list for capture-seq

Table S2. All variants discovered in capture-seq.

Table S3. 23 SMAD3 LD variants within the longevity-associated haplotype.

## Notes

### Competing Interest Statement

The authors have declared no competing interest.

